## Supplementary figures for "BET family members Bdf1/2 modulate global transcription initiation and elongation in *Saccharomyces cerevisiae*"

#### Figures S1 – S7

#### Tables S1 – S5

**Table S1.** Yeast strains used in this study.

**Table S2.** Summary of 4-thioU RNA-seq experiments. Related to Figures 1, 3, S1 and S4.

**Table S3.** Results of gene ontology analysis on 25% most BET sensitive genes in yeast (this work) and human (Winter et al., 2017) system. Related to Figure S2.

**Table S4.** Summary of ChEC-seq and ChIP-seq experiments. Related to Figures 2, 3, 4, 5, S3, S4, S5 and S6.

**Table S5.** Summary of ChIP-seq experiments on Rpb1, Ctk1, Bur1 and Spt5. Related to Figures 6 and S7.

**Table S6.** Number of biological replicates collected for all experiments in this study.

### Figure S1

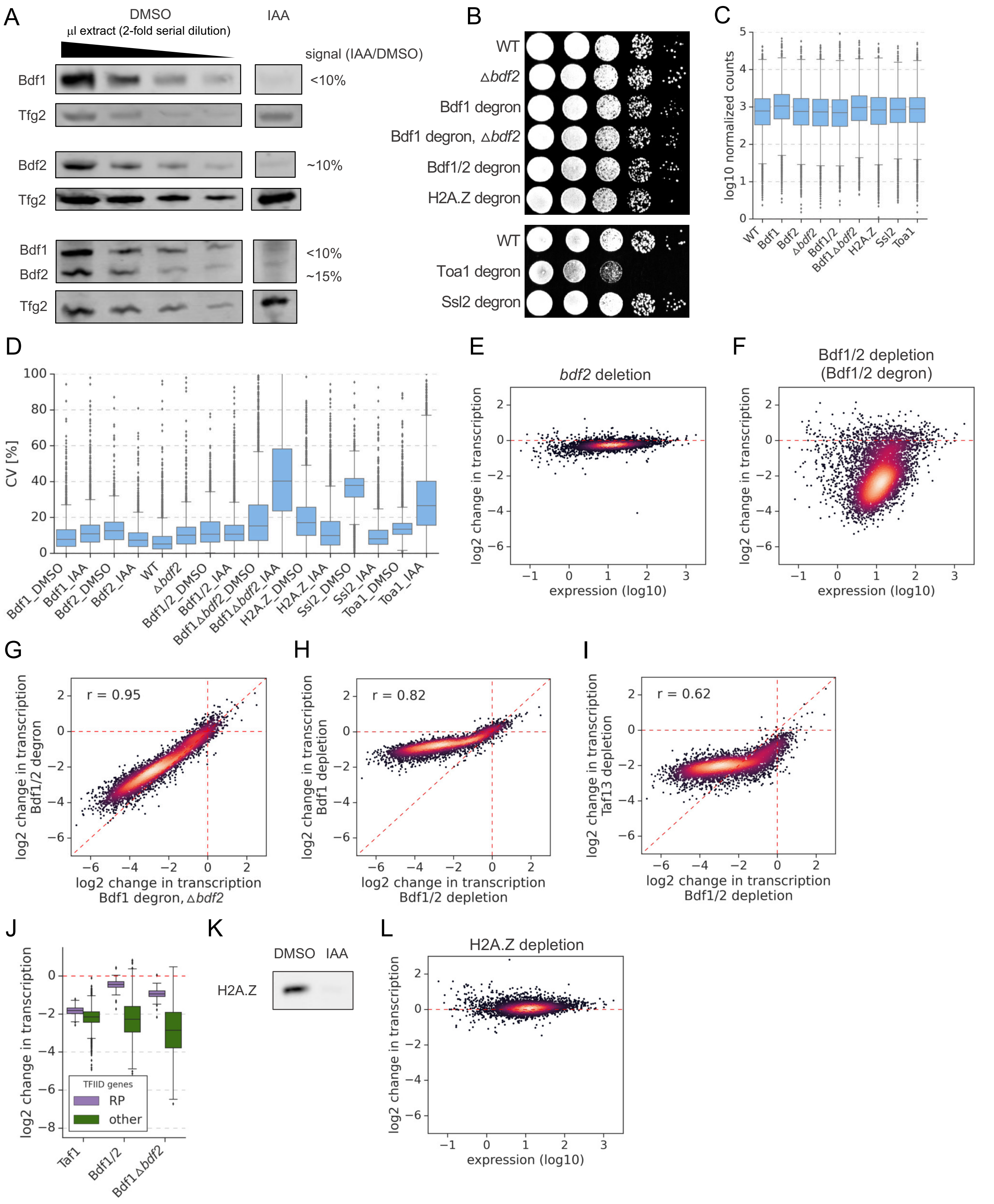
A
DMSO
IAA
µl extract (2-fold serial dilution)
signal (IAA/DMSO)
Bdf1
Tfg2
Bdf2
Bdf1
Bdf2
<10%
~10%
~15%
B
WT
Δbdf2
Bdf1 degron
Bdf1 degron, Δbdf2
Bdf1/2 degron
H2A.Z degron
Toa1 degron
Ssl2 degron
C
log10 normalized counts
D
CV [%]
Bdf1_DMSO
Bdf1_IAA
Bdf2_DMSO
Bdf2_IAA
WT
Δbdf2
Bdf1/2_DMSO
Bdf1/2_IAA
Bdf1Δbdf2_DMSO
Bdf1Δbdf2_IAA
H2A.Z_DMSO
H2A.Z_IAA
Ssl2_DMSO
Ssl2_IAA
Toa1_DMSO
Toa1_IAA
E
bdf2 deletion
log2 change in transcription
expression (log10)
F
Bdf1/2 depletion
(Bdf1/2 degron)
G
r = 0.95
log2 change in transcription Bdf1/2 degron
log2 change in transcription Bdf1 degron, Δbdf2
H
r = 0.82
log2 change in transcription Bdf1 depletion
log2 change in transcription Bdf1/2 depletion
I
r = 0.62
log2 change in transcription Taf13 depletion
log2 change in transcription Bdf1/2 depletion
J
TFIID genes
RP
other
Taf1
Bdf1/2
Bdf1Δbdf2
K
DMSO
IAA
H2A.Z
L
H2A.Z depletion

**Figure S1** (related to Figure 1). (**A**) Western blot analysis of degradation of Bdf1 and/or Bdf2 after 30 min of IAA treatment. Variable amounts of the untreated samples were loaded to allow quantitation of degron efficiency. Blots were probed with $\alpha$-V5 monoclonal (Bdf1, Bdf2) or control $\alpha$-Tfg2 polyclonal antibodies. (**B**) Spot assay comparing the fitness of indicated strains grown at 30°C. A 10-fold serial dilution was used. (**C**) Boxplot of average gene expression level in indicated experiments following DMSO treatment. $Log_{10}$ scale is used for the Y axis. (**D**) Boxplot showing coefficients of variation (CV) for all 5313 genes with detectable signals in all 4-thioU RNA-seq experiments. Samples were collected in two (Bdf1, Bdf2, Δ*bdf2*, Bdf1Δ*bdf2*, H2A.Z) or three (Bdf1/2, Ssl2, Toa1) biological replicates. CV values were calculated for each gene based on normalized read counts for replicate experiments. (**E, F**) Scatter plots comparing expression level (in log scale) with $log_2$ change in transcription per gene in indicated 4-thioU RNA-seq experiments. Mean values from replicate experiments for 5313 genes with detectable signals in all RNA-seq samples collected in this work are plotted. (**G**) Scatter plot comparing $log_2$ change in transcription after depleting Bdf1/2 using two different approaches as indicated. Spearman correlation coefficient (*r*) is shown. Data for 4883 genes classified into TFIID-dependent and coactivator-redundant (CR) categories are plotted (Donczew et al., 2020). (**H**) Scatter plot comparing $log_2$ change in transcription at 4883 genes after depleting Bdf1/2 or Bdf1. Spearman correlation coefficient (*r*) is shown. (**I**) Scatter plot comparing $log_2$ change in transcription at 4883 genes after depleting Bdf1/2 or Taf13. Spearman correlation coefficient (*r*) is shown. (**J**) Boxplot comparing $log_2$ change in transcription at ribosomal protein (RP) genes with the rest of TFIID-dependent genes in the indicated degron experiments. (**K**) Western blot analysis of IAA induced degradation of H2A.Z. Blot was probed with $\alpha$-V5 monoclonal antibody. (**L**) Scatter plot comparing expression level (in log scale) with $log_2$ change in transcription per gene after depleting H2A.Z. Data are shown for 5313 genes with detectable signals in all RNA-seq samples collected in this work.

### Figure S2

A

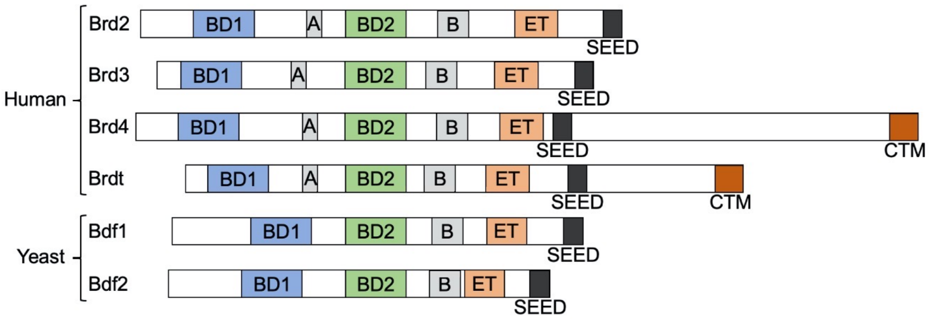


B

| | Bdf1 BD1 | Bdf1 BD2 | Brd4 BD1 | Brd4 BD2 | Taf1 BD1 | Taf1 BD2 |
|---|---|---|---|---|---|---|
| **Bdf1 BD1** | | 1630 | 1570 | 1400 | 920 | 1080 |
| **Bdf1 BD2** | 1630 | | 1800 | 2560 | 820 | 800 |
| **Brd4 BD1** | 1570 | 1800 | | 2110 | 950 | 1320 |
| **Brd4 BD2** | 1400 | 2560 | 2110 | | 1160 | 1190 |
| **Taf1 BD1** | 920 | 820 | 950 | 1160 | | 2500 |
| **Taf1 BD2** | 1080 | 800 | 1320 | 1190 | 2500 | |

C

| Common | Human specific |
|---|---|
| DNA repair | NF-kappaB signaling pathway |
| cell cycle | mitochondrion organization |
| TOR signaling pathway | Pol II transcription |
| autophagy | |
| RNA processing | |
| cellular transport | |
| protein modification | |

**Figure S2** (related to Figure 1). (**A**) Conserved domain organization of human and yeast BET family members. Adapted from (Wu and Chiang, 2007). (**B**) Pairwise alignment scores between amino acid sequences of individual bromodomains of Bdf1, Brd4 and human Taf1 obtained using Clustal Omega (Sievers et al., 2011). (**C**) Major gene ontology terms enriched among the 25% most BET sensitive genes in yeast and human cells.

### Figure S3

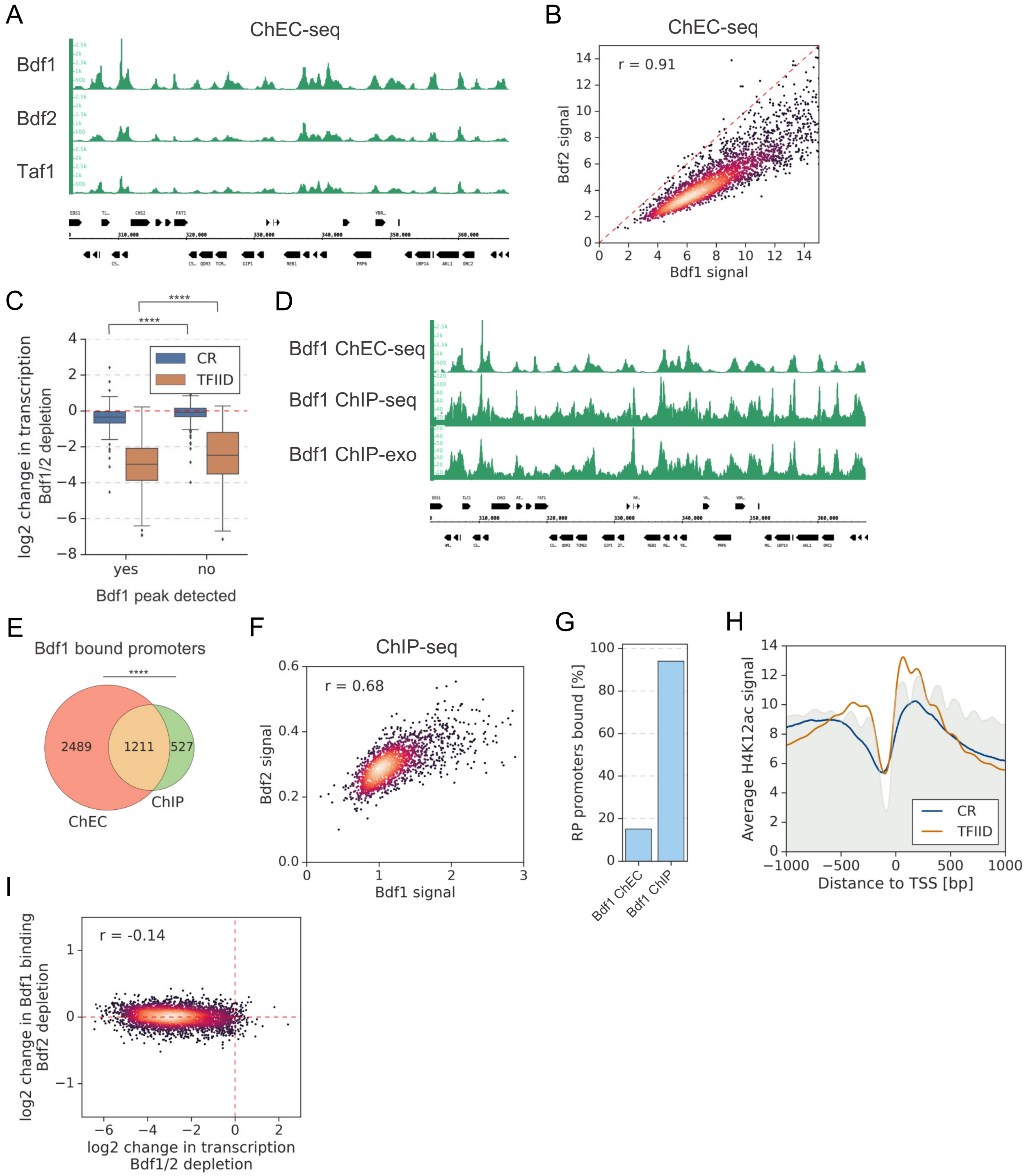

A
ChEC-seq
Bdf1
Bdf2
Taf1
B
ChEC-seq
r = 0.91
Bdf2 signal
Bdf1 signal
C
log2 change in transcription Bdf1/2 depletion
CR
TFIID
yes
no
Bdf1 peak detected
D
Bdf1 ChEC-seq
Bdf1 ChIP-seq
Bdf1 ChIP-exo
E
Bdf1 bound promoters
2489
1211
527
ChEC
ChIP
F
ChIP-seq
r = 0.68
Bdf2 signal
Bdf1 signal
G
RP promoters bound [%]
Bdf1 ChEC
Bdf1 ChIP
H
Average H4K12ac signal
CR
TFIID
Distance to TSS [bp]
I
r = -0.14
log2 change in Bdf1 binding Bdf2 depletion
log2 change in transcription Bdf1/2 depletion

**Figure S3** (related to Figure 2). (**A**) Genome browser image showing comparison of Bdf1, Bdf2 and Taf1 ChEC-seq signals at a representative genomic location. (**B**) Scatter plot comparing Bdf1 and Bdf2 ChEC-seq signals at 3294 promoters bound by both factors. Spearman correlation coefficient (*r*) is shown. (**C**) Boxplot showing the $\log_2$ change in transcription for 4883 genes after depleting Bdf1/2. Genes are grouped into TFIID-dependent and CR categories and data are plotted separately depending on the presence of a significant Bdf1 peak as detected by ChEC-seq. Results of the Welch's t-test are shown. (**D**) Genome browser image showing comparison of Bdf1 ChEC-seq, ChIP-seq (this work) and ChIP-exo signal (Rhee and Pugh, 2012). The same genomic location as in Figure S3A is shown. (**E**) Venn diagram showing the overlap of promoters bound by Bdf1 as detected by ChEC-seq and ChIP-seq. Results of the hypergeometric test are shown. The asterisks represent p-value with the following cut-off levels: **** - 0.0001, *** - 0.001, ** - 0.01, * - 0.05, ns - >0.05. (**F**) Scatter plot comparing Bdf1 and Bdf2 ChIP-seq signals at individual promoters bound by both factors. Spearman correlation coefficient (*r*) is shown. 1738 Bdf1 bound promoters identified by ChIP-seq were used for this analysis. Signals were calculated in a -100 to 200 bp window relative to TSS. (**G**) Barplot showing the percentage of ribosomal protein (RP) gene promoters bound by Bdf1 as detected by ChEC-seq or ChIP-seq. (**H**) Average plot of H4K12ac ChIP-seq signal at 4900 TFIID-dependent or CR promoters. Published MNase-seq dataset is shown as a grey area plot (Oberbeckmann et al., 2019). (**I**) Scatter plot comparing $\log_2$ change in transcription and $\log_2$ change in Bdf1 ChEC-seq occupancy after depleting Bdf1/2 or Bdf2, respectively. Spearman correlation coefficient (*r*) is shown. Bdf1 signal was calculated in a 200 bp window centered on a dominant peak assigned to 3426 promoters bound by Bdf1 and classified into TFIID-dependent and CR categories.

### Figure S4

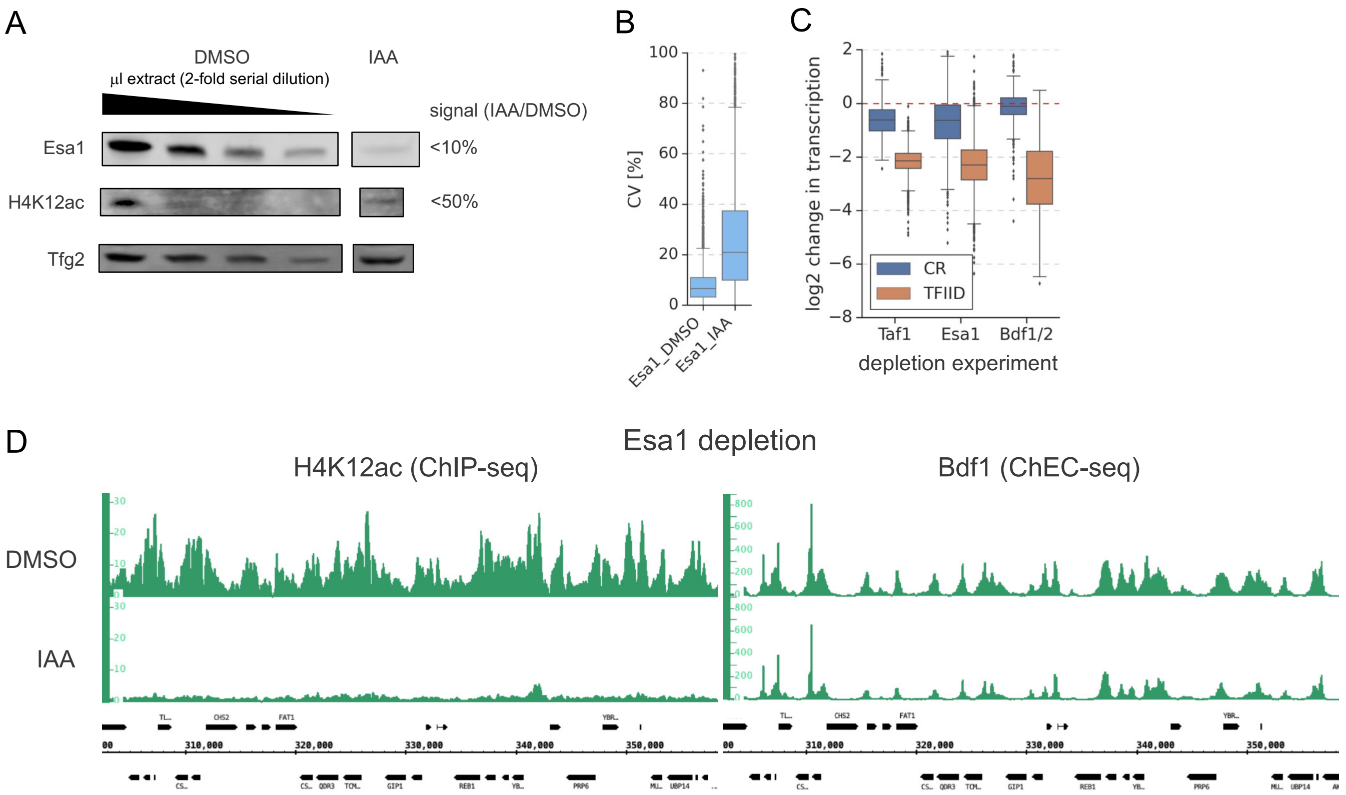


**Figure S4** (related to Figure 3). (**A**) Western blot analysis of degradation of Esa1 and a corresponding change in H4K12ac levels after 60 min of IAA treatment. Variable amounts of the untreated samples were loaded to allow quantitation. Blots were probed with α-V5 monoclonal (Esa1), α-H4K12ac monoclonal and control α-Tfg2 polyclonal antibodies. (**B**) Boxplot showing coefficients of variation (CV) for the Esa1 4-thioU RNA-seq experiment. Two biological replicates were collected. CV values were calculated for each gene based on normalized read counts for replicate experiments. (**C**) Boxplot showing $log_2$ change in transcription at 4883 genes in indicated 4-thioU RNA-seq experiments. Genes are grouped into TFIID-dependent and CR categories. (**D**) Genome browser images showing H4K12ac ChIP-seq and Bdf1 ChEC-seq signals at a representative genomic location before (DMSO) and after (IAA) Esa1 depletion.

### Figure S5

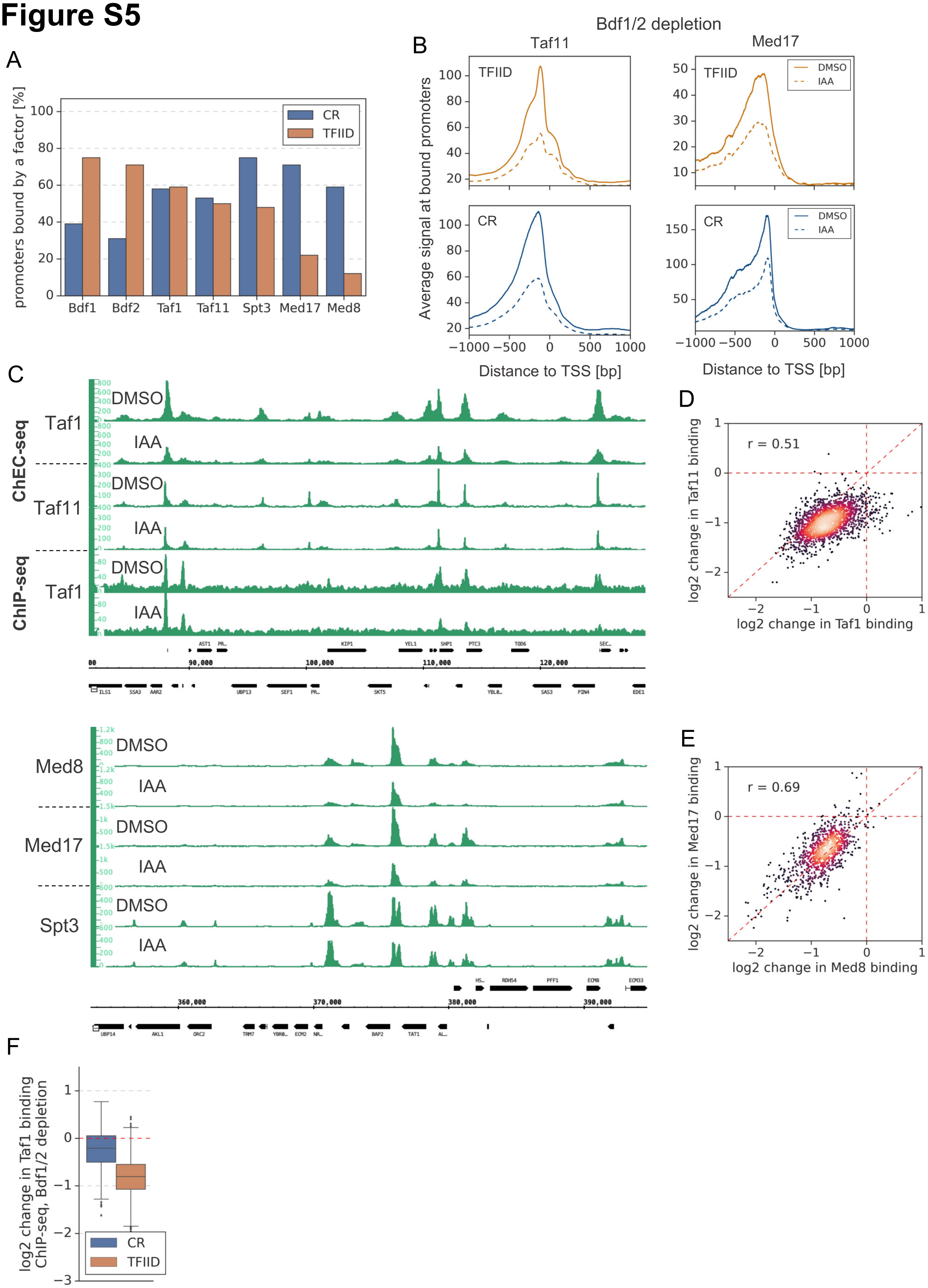

A
promoters bound by a factor [%]
CR
TFIID
Bdf1
Bdf2
Taf1
Taf11
Spt3
Med17
Med8
B
Bdf1/2 depletion
Taf11
Med17
TFIID
CR
DMSO
IAA
Average signal at bound promoters
Distance to TSS [bp]
C
ChEC-seq
ChIP-seq
Taf1
Taf11
DMSO
IAA
Med8
Med17
Spt3
D
r = 0.51
log2 change in Taf11 binding
log2 change in Taf1 binding
E
r = 0.69
log2 change in Med17 binding
log2 change in Med8 binding
F
log2 change in Taf1 binding ChIP-seq, Bdf1/2 depletion
CR
TFIID

**Figure S5** (related to Figure 4). (**A**) Bar plot showing percentage of promoters in each class (TFIID-dependent or CR) bound by a given factor. (**B**) Average plots comparing Taf11 and Med17 ChEC-seq signals before (DMSO, solid line) and after (IAA, dashed line) Bdf1/2 depletion at promoters bound by each factor and classified into TFIID-dependent and CR categories (2486 and 1414, respectively). Mean values from replicate experiments are plotted. (**C**) Genome browser images showing Taf1 ChEC, Taf11 ChEC and Taf1 ChIP (upper panel) or Med8 ChEC, Med17 ChEC and Spt3 ChEC (lower panel) signals at representative genomic locations before (DMSO) and after (IAA) Bdf1/2 depletion. (**D**) Scatter plot comparing $\log_2$ change in binding of Taf1 and Taf11 after depleting Bdf1/2 at 2070 promoters bound by both Taf1 and Taf11 and classified into TFIID-dependent and CR categories. Spearman correlation coefficient (*r*) is shown. (**E**) Scatter plot comparing $\log_2$ change in binding of Med8 and Med17 after depleting Bdf1/2 at 853 promoters bound by both Med8 and Med17 and classified into TFIID-dependent and CR categories. Spearman correlation coefficient (*r*) is shown. (**F**) Boxplot showing $\log_2$ change in ChIP-seq promoter occupancy of Taf1 after Bdf1/2 depletion. Signals were calculated in a -100 to 200 bp window relative to TSS at 2879 promoters defined as bound by Taf1 in ChEC-seq experiments and classified into TFIID-dependent and CR categories.

### Figure S6

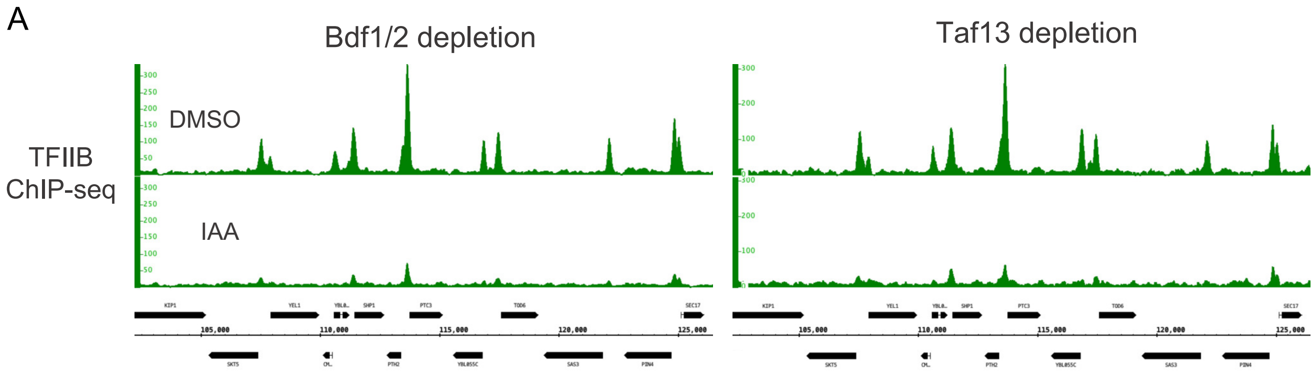


**Figure S6** (related to Figure 5). (**A**) Genome browser images showing TFIIB ChIP-seq signals at a representative genomic location before (DMSO) and after (IAA) Bdf1/2 or Taf13 degradation.

### Figure S7

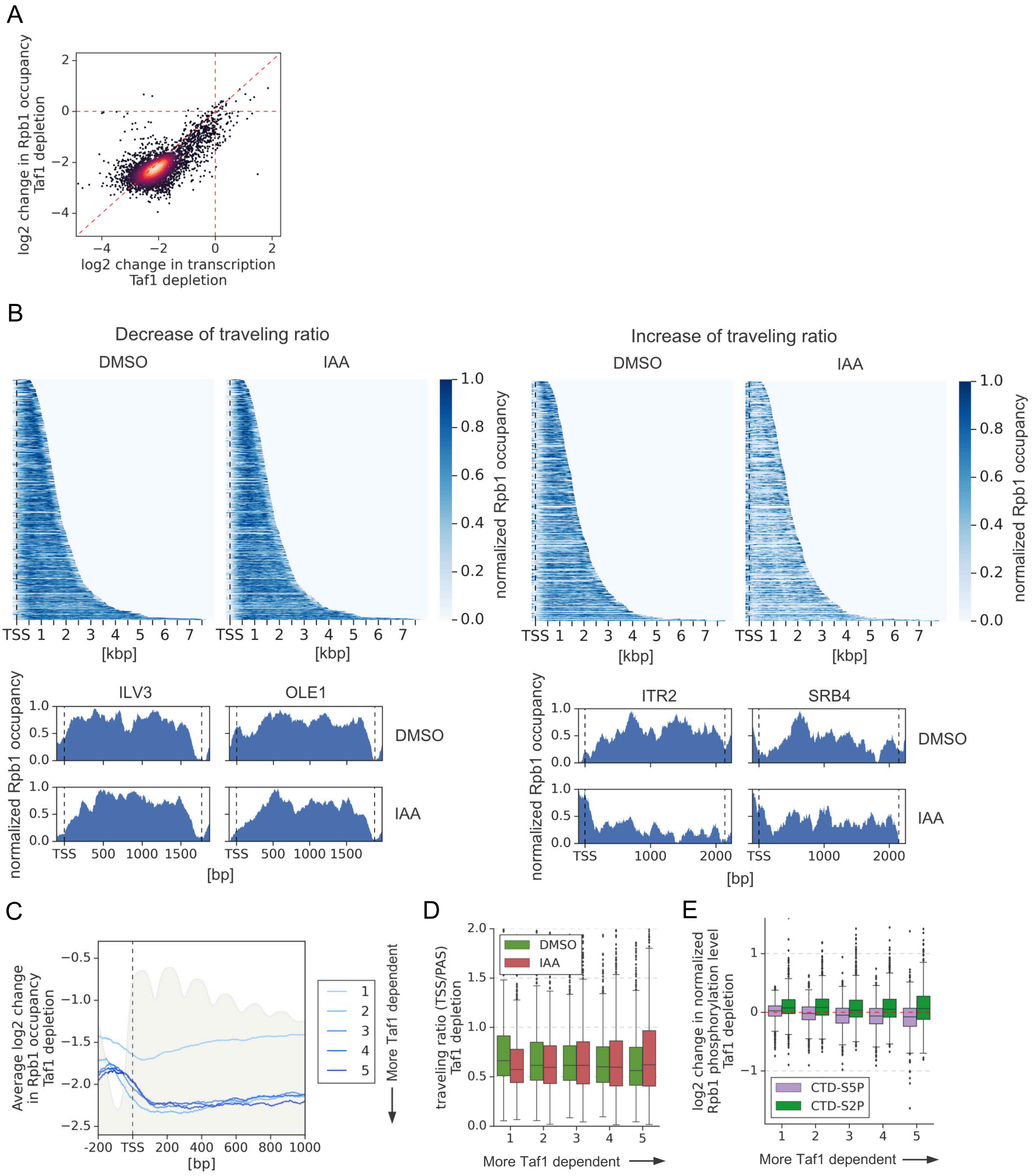
A
log2 change in Rpb1 occupancy Taf1 depletion
log2 change in transcription Taf1 depletion
B
Decrease of traveling ratio
Increase of traveling ratio
DMSO
IAA
normalized Rpb1 occupancy
TSS 1 2 3 4 5 6 7
[kbp]
ILV3
OLE1
ITR2
SRB4
[bp]
C
Average log2 change in Rpb1 occupancy Taf1 depletion
More Taf1 dependent
D
traveling ratio (TSS/PAS) Taf1 depletion
More Taf1 dependent
E
log2 change in normalized Rpb1 phosphorylation level Taf1 depletion
CTD-S5P
CTD-S2P
More Taf1 dependent

**Figure S7** (related to Figure 6). (**A**) Scatter plot comparing $log_2$ change in transcription and occupancy of the largest Pol II subunit Rpb1 after Taf1 degradation. Rpb1 occupancy was calculated along the whole transcribed region for 4615 genes longer than 300 bp and with annotated TSS and PAS locations (Park et al., 2014)). Mean values from replicate experiments are plotted. (**B**) Upper panel: Heatmap showing Rpb1 occupancy at 10% (462) genes with the biggest decrease (left) and 10% (462) genes with the biggest increase (right) in Pol II traveling ratio after depleting Bdf1/2. Data were normalized individually for each gene. TSS position is marked with a dashed line. Separate plots for DMSO and IAA treated samples are shown. Lower panel: Examples of Rpb1 distribution at representative genes with the biggest decrease (left) and the biggest increase (right) in Pol II traveling ratio after depleting Bdf1/2. TSS and PAS positions are marked with dashed lines. Data were normalized individually for each gene. Separate plots for DMSO and IAA treated samples are shown. (**C**) Average plot showing $log_2$ change in Rpb1 occupancy from 200 bp upstream to 1000 bp downstream of TSS after depleting Taf1. Data in this and following panels are divided into five groups based on gene dependence on Taf1 as measured by 4-thioU RNA-seq. 3438 genes longer than 1 kb and with annotated TSS and PAS locations were used (Park et al., 2014). Published MNase-seq dataset is shown as a grey area plot (Oberbeckmann et al., 2019). (**D**) Boxplot comparing Pol II traveling ratio (TR) in DMSO or IAA treated samples in Taf1 degron experiment. TR represents a ratio of Rpb1 occupancy along a 100 bp window at the beginning and end of a transcribed region. The same set of 4615 genes as in Figure S7A was used. (**E**) Boxplot comparing $log_2$ change in Rpb1 CTD phosphorylation status at Ser2 and Ser5 residues after depleting Taf1. Data were calculated along the whole transcribed region for the same set of 4615 genes as in Figure S7A.
